# Permutation-based group sequential analyses for cognitive neuroscience

**DOI:** 10.1101/2023.02.27.530244

**Authors:** John P. Veillette, Letitia Ho, Howard C. Nusbaum

## Abstract

Cognitive neuroscientists have been grappling with two related experimental design problems. First, the complexity of neuroimaging data (e.g. often hundreds of thousands of correlated measurements) and analysis pipelines demands bespoke, non-parametric statistical tests for valid inference, and these tests often lack an agreed-upon method for performing a priori power analyses. Thus, sample size determination for neuroimaging studies is often arbitrary or inferred from other putatively but questionably similar studies, which can result in underpowered designs – undermining the efficacy of neuroimaging research. Second, when meta-analyses estimate the sample sizes required to obtain reasonable statistical power, estimated sample sizes can be prohibitively large given the resource constraints of many labs. We propose the use of sequential analyses to partially address both of these problems. Sequential study designs – in which the data is analyzed at interim points during data collection and data collection can be stopped if the planned test statistic satisfies a stopping rule specified a priori – are common in the clinical trial literature, due to the efficiency gains they afford over fixed-sample designs. However, the corrections used to control false positive rates in existing approaches to sequential testing rely on parametric assumptions that are often violated in neuroimaging settings. We introduce a general permutation scheme that allows sequential designs to be used with arbitrary test statistics. By simulation, we show that this scheme controls the false positive rate across multiple interim analyses. Then, performing power analyses for seven evoked response effects seen in the EEG literature, we show that this sequential analysis approach can substantially outperform fixed-sample approaches (i.e. require fewer subjects, on average, to detect a true effect) when study designs are sufficiently well-powered. To facilitate the adoption of this methodology, we provide a Python package “niseq” with sequential implementations of common tests used for neuroimaging: cluster-based permutation tests, threshold-free cluster enhancement, *t*-max, *F*-max, and the network-based statistic with tutorial examples using EEG and fMRI data.

## 1. Introduction

Recently, many scientific fields have been placing a renewed emphasis on issues of sample size determination and statistical power. This emphasis is motivated, in large part, by an increased appreciation of the fact that the positive predictive value of a study – that is, the probability that an effect is actually “true” given a statistically significant result – is directly proportional to the statistical power of the study (Ioannidis, 2005). This methodological concern came into focus in the neurosciences after a landmark review in 2013 estimated that the median statistical power of neuroscience studies is between 8% and 31%, suggesting that many neuroscience studies provide low evidentiary value despite satisfying conventional standards of statistical evidence (Button et al., 2013).

The average statistical power of neuroimaging studies, in particular, has been steadily improving; however, low statistical power is cited as one of the largest threats to the replicability of findings in cognitive neuroscience (Poldrack et al., 2017). The availability of large, open datasets such as the Human Connectome Project (Van Essen et al., 2013) and tools for extracting metadata from published neuroimaging studies such as NeuroSynth (Yarkoni et al., 2011) have enabled empirical estimates of power in the field. As a result, we now know that the effect sizes one can realistically expect in neuroimaging studies are usually quite small, and many published neuroimaging studies are too small to detect them (Poldrack et al., 2017). Indeed, a recent analysis has suggested that certain types of neuroimaging biomarkers may even require *thousands* of subjects to detect reliably (Marek et al., 2022), though other researchers have been quick to point out that not all analytic approaches require such prohibitive sample sizes to establish robust brain-behavior relationships (Rosenberg & Finn, 2022). In any event, the costs of doing well-powered neuroimaging research can be substantial.

In light of these field-wide concerns, it is increasingly acknowledged that researchers should determine their sample-size in a rigorous, non-arbitrary manner; the use of heuristics, such as adapting the same sample size as a previous study, is likely to result in an underpowered design (Poldrack et al., 2017). Some efforts – notably Neuropower, Fmripower, and PowerMap – have provided researchers with the tools to perform power analyses for the parametric random-field theory approaches used in fMRI (Durnez et al., 2016; Joyce & Hayasaka, 2012; Mumford & Nichols, 2008). However, these tools remain limited relative to the scope of statistical tests employed in the broader neuroimaging literature. In particular, many of the statistical tests used in neuroimaging are non-parametric, and thus parametric power analysis procedures are inapplicable (though a power analysis may be performed by simulation or resampling). Moreover, even in cases in which a validated method for performing a power analysis would be straightforward, specifying an “effect size” is not as simple as for univariate tests, in which power analyses are performed with standard effect size measures (e.g. Cohen’s *d);* neuroimaging analyses are often performed on aggregate properties of spatiotemporal maps, not necessarily on specific voxels. Indeed, an effect map (which would be difficult to predict a priori) is often precisely what the researcher is trying to estimate when they are designing their experiment. Moreover, even in the univariate case, obtaining a reasonably precise effect size estimate usually requires a larger sample size than that required to merely detect an effect (Albers & Lakens, 2018; Lakens & Evers, 2014), and it understandably belies most researchers’ intuition to collect a pilot sample larger than their confirmatory study.

Most published fMRI and EEG studies still do not include a sample size justification, likely due to the substantial challenges associated with performing a power analysis described above. For instance, out of 100 clinical fMRI studies randomly sampled from six leading journals, only a single study reported a sample size calculation (Guo et al., 2014). Similarly, 0 out of 100 randomly sampled studies from the EEG literature reported sample size calculations (Larson & Carbine, 2017). We do not believe that this omission results from researchers’ lack of desire to do more rigorous science, but rather it is the result of a lack of methodological approaches and tools that meet their specific research needs. Indeed, when reviewers for granting agencies such as NSF and NIH request sample size calculations for proposed research, PIs may scramble to find a way of estimating these, whether rigorous or not.

One alternative researchers may find tempting is to forgo an a priori sample size determination and analyze data multiple times throughout the course of data collection, stopping data collection only once a significant result is found. Indeed, many researchers in psychology admit to employing this practice known as *optional stopping* (John et al., 2012). However, optional stopping results in inflated false-positive rates; for instance, if one analyzes the data five times throughout data collection without adjusting their significance threshold of *α* = 0.05, the false positive rate rises to an undesired 0.142 (Armitage et al., 1969). Again, however, we do not believe researchers admit to optional stopping because of a negligible disregard for research best-practices, but because of a desire to preserve time and resources, stopping data collection as soon as there is sufficient evidence to reach a conclusion. Indeed, one could argue researchers have an ethical obligation to use (often taxpayer-funded) resources efficiently and to limit the burden on the human subject populations that volunteer for their experiments.

Fortunately, valid *sequential analysis* techniques, which use adjusted rejection thresholds at each interim analysis to control the false positive rate across the whole experiment, have existed for the better part of a century (Dodge & Romig, 1929; Wald, 1992). Sequential study designs have long been recognized to possess efficiency advantages over their sequential counterparts, allowing a conclusion to be reached with fewer observations on average (Dodge & Romig, 1929). The reason for this efficiency advantage is straightforward. For instance, if one analyzes the data once at a sample size where there is a 50% chance of rejecting the null hypothesis and once at *n*_2_ when there is a 95% chance of rejection, then the expected sample size at which a conclusion is reached would likely end up somewhere between and *n*_2_, since the probability of rejection at *n*_1_ is substantial.

While early approaches to sequential analysis required data to be analyzed after every observation or, alternatively, the exact number and timing of looks at the data to be specified a priori (a *group* sequential analysis), which was somewhat limiting for studies where the final subject yield may not be known until the data are analyzed (e.g. after quality check of neuroimaging data), a later approach known as *alpha spending* requires only that a maximum sample size (that is, the sample size at which data collection will be terminated even if a significant result has not been reached) be specified ahead of time (G. Lan & DeMets, 1983). This approach is heavily used in the clinical trial literature and has recently attracted attention as a means of improving the efficiency of experimental psychology studies as well (Lakens, 2014). While we believe the general alpha spending approach is sufficiently flexible to meet the practical demands of neuroimaging studies, the adjusted significance thresholds it prescribes for interim analyses are only valid assuming test statistics across looks follow a multivariate normal distribution (G. Lan & DeMets, 1983), an assumption that is violated by many of the test statistics in neuroimaging.

To this end, we propose a permutation-based version of the alpha spending procedure. This procedure allows sequential analyses to be performed using arbitrary test statistics, and we show by simulation that it controls the false positive rate while doing so. Since many hypothesis tests and multiple-comparisons corrections used in cognitive neuroscience are already permutation-based, they can be naturally generalized into valid sequential tests. Thus, we were able to implement sequential generalizations of cluster-based permutation tests (Maris & Oostenveld, 2007), threshold-free cluster enhancement (Smith & Nichols, 2009), *t*-max and *F*-max corrections (Nichols & Holmes, 2002), and the network-based statistic (Zalesky et al., 2010). As such, the proposed approach to sequential-testing can be easily applied to task EEG and fMRI, functional connectivity, diffusion tractography, and voxel-based morphometry data.

This sequential testing approach affords several routes to principled sample size determination that were previously unavailable to human neuroscience researchers. (1) One approach is to specify a very conservative maximum sample size; if the maximum sample is overly conservative, one will reject the null hypothesis at an interim analysis with high probability. (2) Another approach is to use an *adaptive design*, in which the maximum sample size can be adjusted as a result of a *conditional power analysis* performed at an interim analysis (Lakens et al., 2021). In this case, the first interim analysis can serve as an internal pilot study, used to estimate the effect size with which a power analysis is performed (which can be done very generally via resampling methods). However, if enough evidence has already been accrued at that interim analysis to reject the null hypothesis, then data collection can be concluded. (3) Lastly, even if a researcher has everything needed to perform an a priori sample size calculation, it may still be advantageous to use a sequential design so that data collection can be terminated early if enough evidence has been accrued to reject the null hypothesis. We show that, like parametric alpha spending, our approach can be substantially more efficient than fixed-sample designs for studies that are sufficiently well-powered.

## 2. Methods

### 2.1. Alpha spending

#### 2.1.1. General description

In an alpha spending procedure, one must decide two things in advance of beginning data collection. (1) One must specify a maximum sample size after which data collection will be stopped, regardless of whether the null hypothesis has been rejected. We will call this design parameter *n*_T_ throughout this article, where *T* denotes the total number of interim analyses. Note, however, that *T* does not need to be determined before the study begins, only *n_τ_* = *n*_max_. (2) Additionally, one must define a desired Type I error rate *α* and an *alpha spending function* that specifies how the Type I error rate will be distributed across interim analyses. Specifically, an alpha spending function *s*(*n*) is a monotonically non-decreasing function that specifies a target value for the *cumulative* Type I error by interim sample size *n*. Consequently, the function *s*’ must also satisfy *s*(0) = 0 and *s*(*n*_T_) = *α* for the desired significance level *α*, such that the Type I error rate is contained to *α* at *n*_T_. Within these constraints, any function *s*: ℝ_1_ → ℝ_1_ (i.e. that maps a real number to a real number) could be specified.

Suppose the data are analyzed at each of the successive interim sample sizes 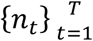. (This notation more concisely denotes a set containing ordered interim sample sizes *n*_2_, …, *n*_2_,…, *n*_*T*-1_, *n_T_* for an arbitrary positive integer *T*.) At each interim sample size *n_i_*, then, the data are 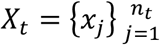 (where the index of *x_i_*, importantly, denotes the order in which the observation was collected rather than being a random index) with condition labels (or covariate of interest) 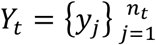, and the test statistic is *F_t_* = *f*(*X_t_, Y_t_*), where *f* is some function that computes the test statistic from the data.

We need to determine some threshold 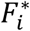 at each interim analysis that will contain the cumulative Type I error, given all the previous looks, to *s*(*n_t_*). That is, for each interim analysis *t* ∈ [1, *T*], we select 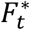 so as to satisfy Equation 1.

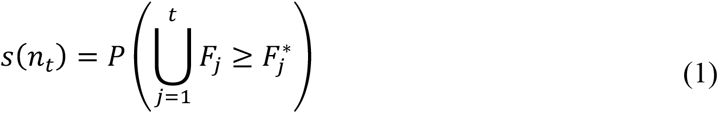

Note that the right side of Equation 1 can be expanded as in Equation 2.

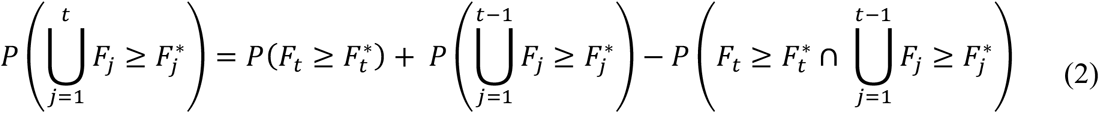

It is this joint probability on the right side of Equation 2 that can be difficult to compute, but it will almost certainly be nonzero since the *F_t_*’s are computed from overlapping data. In the parametric approach to alpha spending introduced by Lan and DeMets, the joint distribution of the test statistics 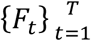 is assumed to be multivariate normal under the null hypothesis, which allows appropriate rejection thresholds 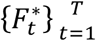 to be computed exactly. In the event that this normality assumption is violated, these thresholds must be estimated in some other way.

Note that, by plugging the rejection thresholds 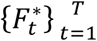 into the cumulative distribution function of *F_t_* under the null hypothesis, it is possible to obtain adjusted significance levels 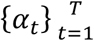 for each look time. It is common to provide these adjusted significance levels when reporting the results of a sequential analysis (Lakens et al., 2021).

#### 2.1.1. Common spending functions

As noted above, there is considerable flexibility in the selection of an alpha spending function *s*(*n*); any function that satisfies the constraints specified in the above section will be adequate. However, it is worth mentioning a few common spending functions used in the literature. Lan and DeMets introduced two spending functions, the Pocock and O’Brien Fleming spending functions (G. Lan & DeMets, 1983). The Pocock spending function “spends” the Type I error rate more liberally early during data collection, so it is more likely to reject the null hypothesis early on if the effect size is large. The O’Brien Fleming spending function, on the other hand, “saves” its Type I error allotment for later during data collection when there is more power to detect a given effect, so while it is less likely to terminate very early if the effect size is unexpectedly large, it may be more likely to reject the null hypothesis at an interim analysis overall. A linear spending function, which falls somewhere between those two, distributed the error rate evenly across data collection. We recommend consulting Lakens and colleagues’ tutorial on group sequential designs for a more thorough discussion and comparison of adjusted alpha thresholds between spending functions (Lakens et al., 2021).

#### 2.1.2. Inflation factors and expected sample size

Generally speaking, if one wants to design a study using alpha spending that has the same statistical power as a given fixed sample design with sample size *n*_fixed_, then the maximum sample size *n_T_* for the sequential design will need to be greater than *n*_fixed_. The ratio *n_T_*/*n*_fixed_ required for the two designs to have matched power is called the *inflation factor*, and it depends on the alpha spending function, the number and timing of interim analyses, and the desired statistical power. However, usefully, it does not depend on *n*_fixed_, the effect size, or the statistical test used.

Even more usefully, the inflation factors for parametric alpha spending designs are known exactly, and they can be easily obtained from open-source statistical software such as rpact (Wassmer & Pahlke, 2020). While these inflation factors are not theoretically guaranteed to apply to the permutation alpha spending approach we describe below, empirically they seem to work quite well (see our simulation results below).

While the inflation factor *n_T_*/*n*_fixed_ needed to obtain matched statistical power may generally be positive (i.e. the maximum sample size for the sequential design must be larger than the fixed-sample design), this inflation is offset by the fact that sequential designs have the opportunity to reject the null hypothesis at an interim analysis and therefore *n_T_* observations may never actually be collected. The *expected sample size* of a sequential design, then, differs from the maximum sample size.

The expected sample size depends on the power of the sequential design. If the study is sufficiently well powered, the probability of detecting the effect early is high and the expected sample size is often substantially lower than the corresponding fixed-sample design’s sample size; if, conversely, the design is very poorly powered, then the experiment will likely be run to completion without detecting the effect and the expected sample size will be close to *n_T_*. The expected sample size of a given sequential design, as a function of statistical power, can be found exactly for parametric alpha spending designs and can be easily obtained from software packages like rpact.

### 2.2. Permutation alpha spending

We propose a permutation-based procedure to infer the co-dependence term in Equation 2 implicitly from the data. Conceptually, this is similar to how permutation-based approaches for controlling the familywise error rate outperform parametric approaches (e.g. Bonferroni correction) in the event that one’s multiple tests are correlated, since the covariance between test statistics is implicitly accounted for because that correlation structure is preserved in the empirical null distribution generated by permutating the data (Nichols & Holmes, 2002). That is, the permutation null is a *joint* null distribution.

Let *H*_0_ be a *K* × *T* matrix, where *K* is the number of permutations we decide to perform. We want to populate the matrix *H*_0_ with the *joint null distribution* of the test statistic across the *T* interim analyses, which we will empirically estimate by permutation.

Then for each permutation *k* ∈ [1, *K*] and each *t* ∈ [1, *T*],

1. Let 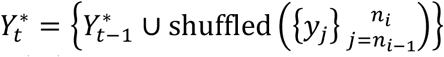, where 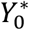 is an empty set.
2. Set 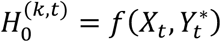

Note that Step 1 above implies that any observation *x_i_* has the same shuffled label at all times *t* ∈ [1, *T*] on a given permutation *k*. Without this feature, *H*_0_ will not be a joint distribution. If one is performing a one-sample test instead of an independent-samples tests, then the signs of the *x_i_*’s are randomly flipped instead of shuffling their *y_i_* labels.

One can then estimate the rejection thresholds from *H*_0_ as follows.

- Pick 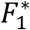 such that only 100 · *s*(*n*_1_) percent of the values in 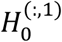 surpass 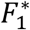.
- In ascending order of *t*, pick each 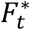 such that only 100 · *s*(*n_t_*) percent of permutations *k* have any 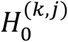 that exceed 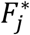 at any *j* ∈ [1, *t*].

That is, each 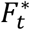 is chosen to control the *cumulative* false positive rate given all previous *F*\* to match the target given by the alpha spending function *s*(*n_t_*). Since all thresholds 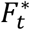 depend on the previous thresholds and data, but not on the later thresholds and data, then rejection thresholds can be computed at the time of each interim analyses *t*, and data collection can be halted if 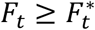.

As in the parametric case, adjusted *α_t_*‘s can be obtained from each 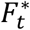 by computing the percentile rank of 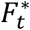 within 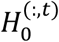, which is approximately equivalent to plugging 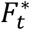 into the empirical cumulative distribution function of *F_t_* under the null hypothesis.

### 2.3. Simulation Studies

#### 2.3.1. False positive rate simulations

First, we validate the permutation scheme by estimating the false positive rate for a univariate test by Monte Carlo simulation. In particular, on each simulation, we generate 500 random variables from the standard Normal distribution. We then compare two approaches. (1) In a simulated optional stopping procedure, we compute a *p*-value using a two-sided, one-sample permutation *t-*test with 1024 permutations at each of *n* = 100,200,300,400,500, and we reject the null hypotheses if *p* ≤ 0.05 at any of these interim analyses. Estimating the false positive rate as the proportion of 10,000 simulations in which the null hypothesis was erroneously rejected, the false positive rate of this optional stopping procedure should be about 0.142 (Armitage et al., 1969). (2) We perform the same interim analyses, but we compute adjusted significance thresholds using the permutation alpha spending procedure described above with a linear spending function. Then, we only reject the null hypothesis if a test statistic exceeds the adjusted threshold at an interim analysis. This procedure should result in a false positive rate of approximately *α* = 0.05.

We additionally perform the above simulations for a between-sample test, in which the simulated observations are randomly assigned to one of two conditions and an independent samples permutation *t*-test is used to compare between conditions.

Lastly, we verify that this procedure still controls the false positive rate for more sophisticated test statistics (e.g. one actually used on effect maps in neuroimaging). This time, we repeat all the above simulations but instead of generating random observations of one variable, we generate observations from 100 variables resulting in 100 different tests at each interim analysis. We control for multiple comparisons at each look time using a *t*-max procedure (Nichols & Holmes, 2002). (1) In the optional stopping procedure, we reject the null hypothesis if a *t*-max adjusted *p*-value (from any of the 100 tests) is less than or equal to 0.05 at any interim analysis. (2) In the sequential analysis procedure, we compute adjusted significance levels using permutation alpha spending with the *t*-max statistic, and we only reject the null hypothesis if a *t*-max adjusted *p*-value is less than or equal to those adjusted thresholds.

#### 2.3.2. Power/efficiency simulations

To assess the efficiency of sequential designs relative to fixed-sample designs, we ran power analyses for detecting seven canonical EEG event-related potential (ERP) effects from the ERP CORE dataset (Kappenman et al., 2021) using both types of design. Specifically, we used the lowpass-filtered versions of the precomputed difference waves for each ERP effect provided in ERP CORE. The dataset contains data from 40 healthy subjects for each ERP effect, though some of the subjects are excluded from certain ERP effects for low quality data. See the ERP CORE reference paper for a full description of data collection and preprocessing (Kappenman et al., 2021).

To estimate the statistical power for a given sample size *n*, we use a resampling technique. Specifically, we use a modified Bayesian bootstrap, which differs from the frequentist bootstrap in that it simulates draws from the posterior distribution (with uninformative priors) of a parameter instead of the sampling distribution of that parameter (Rubin, 1981). Strictly speaking, the frequentist bootstrap is a special case of the Bayesian bootstrap in which the original observations are resampled with the same probability on every bootstrap resampling; the Bayesian bootstrap draws the new resampling probabilities from a Dirichlet distribution on each bootstrap resampling. The result is that “the Bayesian bootstrap can be thought of as a smoothed version of the Efron bootstrap” (Lancaster, 2003), generally yielding more stable results when resampling from smaller pools of observations – though it asymptotically converges with the frequentist bootstrap. Estimating power by Bayesian bootstrap lends itself to interpretation as a non-parametric Bayesian predictive power (Spiegelhalter et al., 1986), which appropriately accounts for uncertainty about the effect size in estimating the power of a frequentist test.

On each simulation (i.e. bootstrap resample), then, we resample *n* observations from the original 40 observations and run our statistical test. Power is estimated as the proportion of simulations in which the null hypothesis is rejected.

For each ERP effect, we estimate the statistical power of a fixed-sample design with *n =* 30. Then, we estimate the maximum sample size *n_τ_* needed to obtain the same power using a sequential design – with a single interim analysis performed midway through data collection and a Pocock spending function – by obtaining the necessary inflation factor (see Section 2.1.2) from rpact (Wassmer & Pahlke, 2020). We then estimate the statistical power of the sequential design in the same manner (i.e. the proportion of simulations in which the null hypothesis is rejected at either the interim or maximum sample size), and we additionally estimate the expected sample size (see Section 2.1.2) as the average of the sample size at which “data collection” was terminated (either because the null hypothesis was rejected or the simulated study was completed without rejecting the null) across all simulations.

Since the ERP CORE consists of highly optimized EEG paradigms, designed to maximally elicit the ERP effect of interest, it is sometimes the case that every subject in the dataset shows the effect individually. If this is the case, our bootstrap procedure might estimate a power of 1, which is not informative for our purposes since an inflation factor cannot be computed for a design of power 1. So, when this occurs, we modify the bootstrap procedure as follows. After drawing our *n* resamples from the original observations on each bootstrap simulation, we generate a *n* noise time series (see next paragraph for noise generation procedure) and add them to the *n* samples. If this addition of noise was not sufficient to decrease power to be less than 1, we ran the simulations again, multiplying the noise by 2. If this failed, we multiplied the noise by 3. No effect required noise be multiplied by a factor greater than 3. The exact noise multipliers used for each effect can be found in our archived results (see Section 2.4: *Data and Code Availability*).

To generate noise, we estimate the covariance (between sensors) matrix of the grand-averaged difference wave for the ERP effect; then, we generate spatially colored multivariate Gaussian white noise using this covariance matrix so as to maximally interfere with the effect of interest. We then lowpass filtered the noise to match the ERP data (which was filtered at 20 Hz), which created temporal autocorrelation in the filtered signal.

10,000 bootstrap simulations were performed as above for each ERP effect using both a *t*-max test and a cluster-based permutation test (with a clustering threshold of *t* = 2), both with 1024 permutations. After running all simulations, (a) we compared the power of the fixed sample designs to that of the sequential designs, which should have a 1:1 relationship if the inflation factors adequately predict the sample size required to match the power of a fixed sample design. (b) Then, we assess the efficiency of the sequential designs by comparing their expected sample sizes to the fixed sample size, which is always 30.

### 2.4. Data and Code Availability

Our implementations of permutation alpha spending for cluster-based permutation tests, threshold-free cluster enhancement, *t*-max, *F*-max, *r*-max, and the network-based statistic are contained in our user-friendly Python package *niseq*, which can be installed from the Python package index (PyPI). Documentation is hosted on Read the Docs (http://niseq.readthedocs.io/). Source code, as well as worked examples using the package on EEG and fMRI data in conjunction with the MNE-Python (Gramfort et al., 2014) and nilearn packages, are available on GitHub (https://github.com/john-veillette/niseq) and permanently archived on Zenodo. The most recent release as of writing (v0.0.2) is available at https://doi.org/10.5281/zenodo.7526535 and the current release is always archived at https://doi.org/10.5281/zenodo.7517285.

The code used for the simulations featured in this article, as well as the results of those simulations and a record of the simulation parameters, are available separately on GitHub (https://github.com/john-veillette/niseq-simulations) and are permanently archived on Zenodo (https://doi.org/10.5281/zenodo.7666443).

The data used for simulations was originally taken from the ERP CORE repository on the Open Science Framework (https://doi.org/10.18115/D5JW4R). However, we exported the preprocessed difference waves into a file format that could be easily loaded with MNE-Python, which we provide for convenience in the same Zenodo archive as our simulation code.

## 3. Results

### 3.1 False positive rates

Permutation alpha spending controls the false positive rate below the specified *α* = 0.05 across multiple interim analyses for single permutation *t-*tests, and it controls the familywise error rate in a sequential *t-*max procedure (see Table 1). In contrast, permutation *t*-tests and *t-*max with optional stopping results in inflated false positive rates.

**Table 1:** False positive rates for optional stopping and for permutation alpha spending procedures with five interim analyses.

| | One sample<br>(univariate) | One sample<br>($t$-max) | Indep. sample<br>(univariate) | Independent<br>($t$-max) |
| --- | --- | --- | --- | --- |
| **Optional stopping** | 0.1442 | 0.1784 | 0.1402 | 0.1715 |
| **Alpha spending** | 0.0470 | 0.0439 | 0.0465 | 0.0458 |

Notably, how to best correct for multiple comparisons in group sequential designs with multiple outcomes of interest is still an active topic of research in the clinical trial literature (Glimm et al., 2010; Kosorok et al., 2004; Tang & Geller, 1999). Sequential *t*-max provides a solution to this problem that, like its fixed-sample counterpart, can scale to hundreds and thousands of arbitrarily correlated tests.

### 3.2 Efficiency relative to fixed-sample designs

The results of our power simulations for fixed-sample and sequential designs are illustrated in Figure 3. When we set the maximum sample size for each sequential design by multiplying the sample size of the fixed-sample design (always *n*_fixed_ = 30) by the appropriate inflation factor, we obtain a design with roughly matched power (see Figure 3a and 3c). In other words, the inflation factors used for parametric sequential designs seem applicable to our approach.

**Figure 1:**
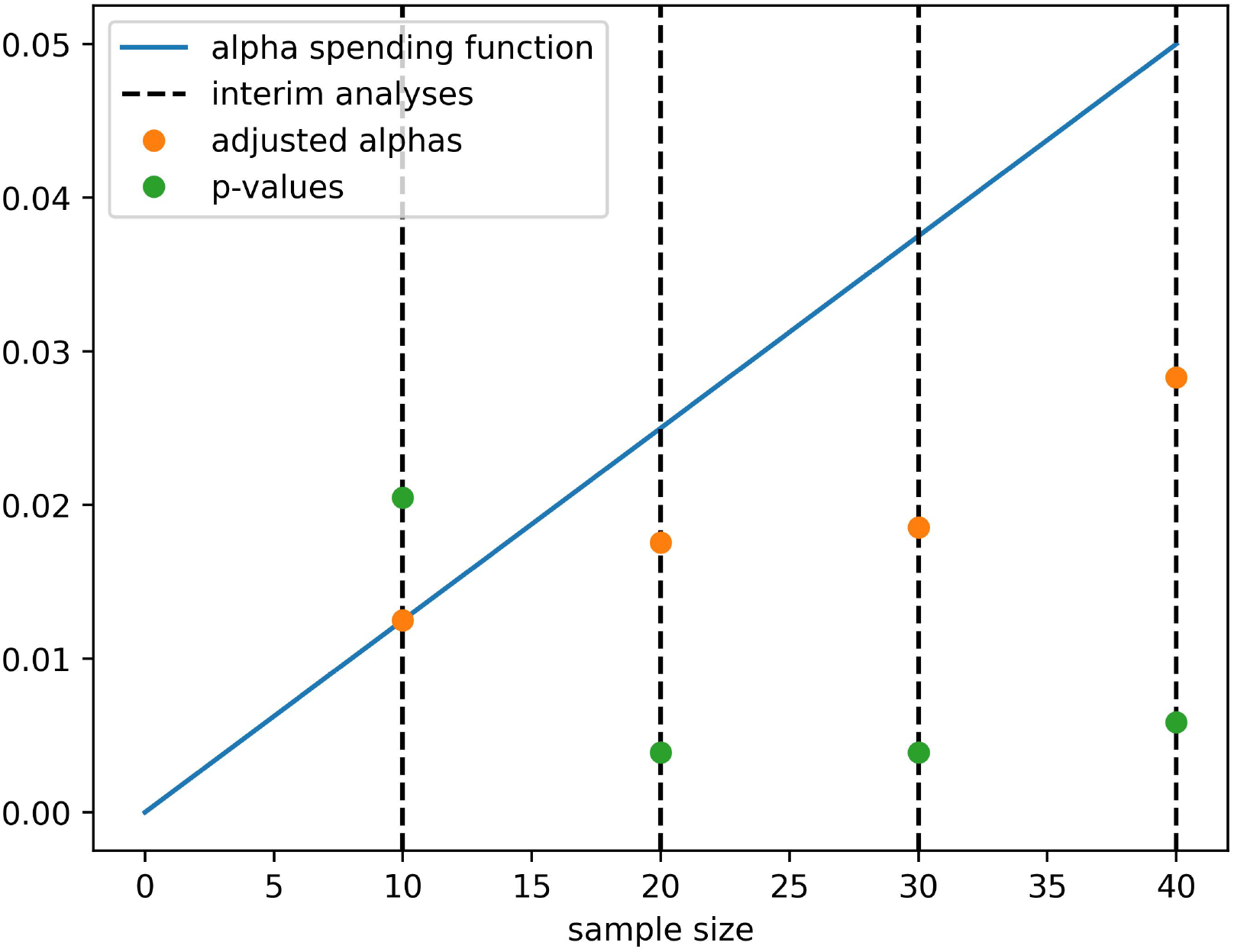
An example of an alpha spending procedure with a linear spending function. At each interim analysis, an adjusted significance level *α* is computed that controls the *cumulative* Type I error, given the previous interim analyses, to the value specified by the alpha spending function. In this example, data collection would be stopped at *n* = 20, where the observed *p*-value drops below the adjusted *α*.

**Figure 2:**
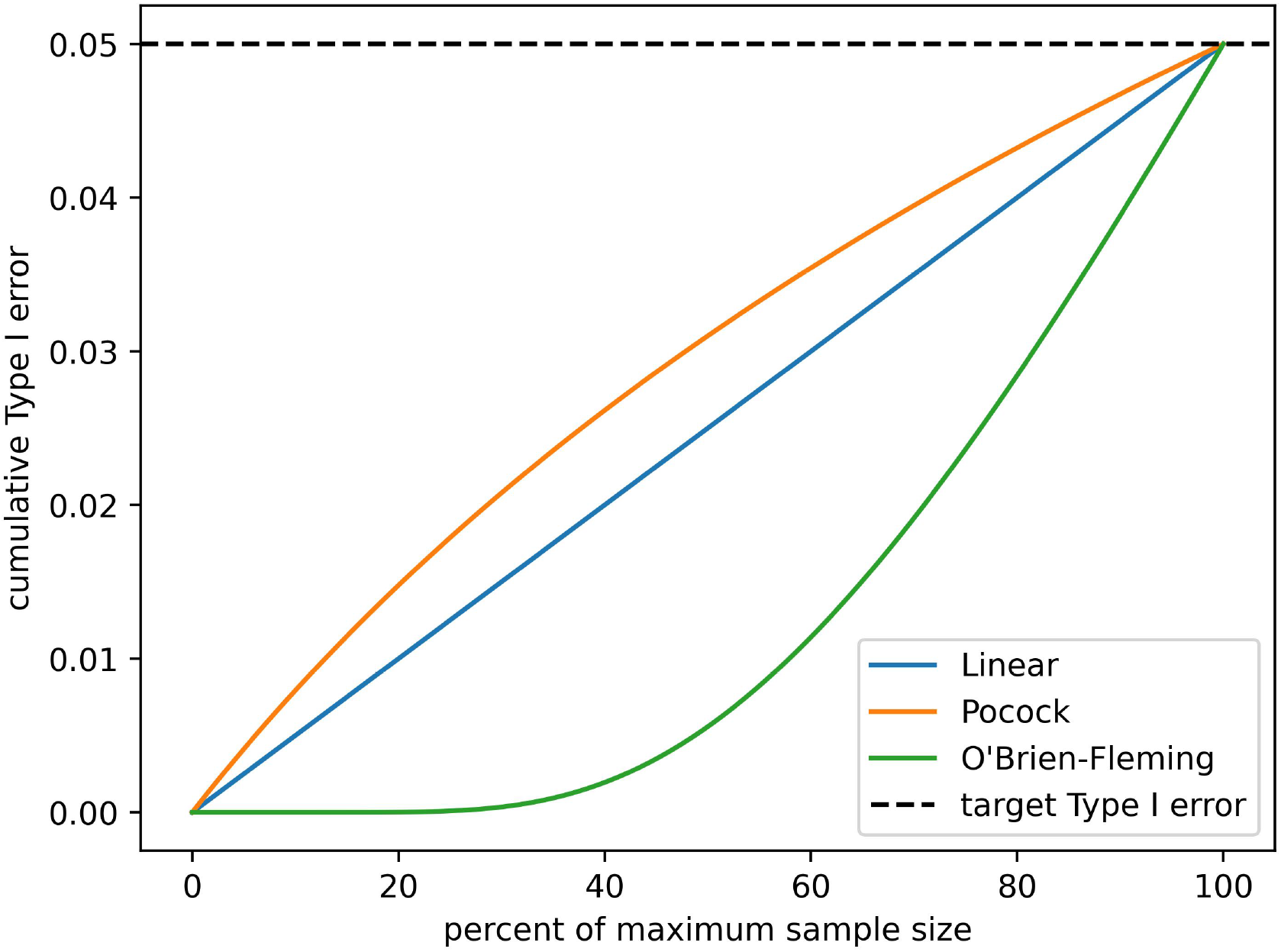
Examples of common alpha spending functions. Spending functions all start at 0 and end at the user-specified false positive (Type I error) rate, but they vary in how they distribute Type I error across interim analyses.

**Figure 3:**
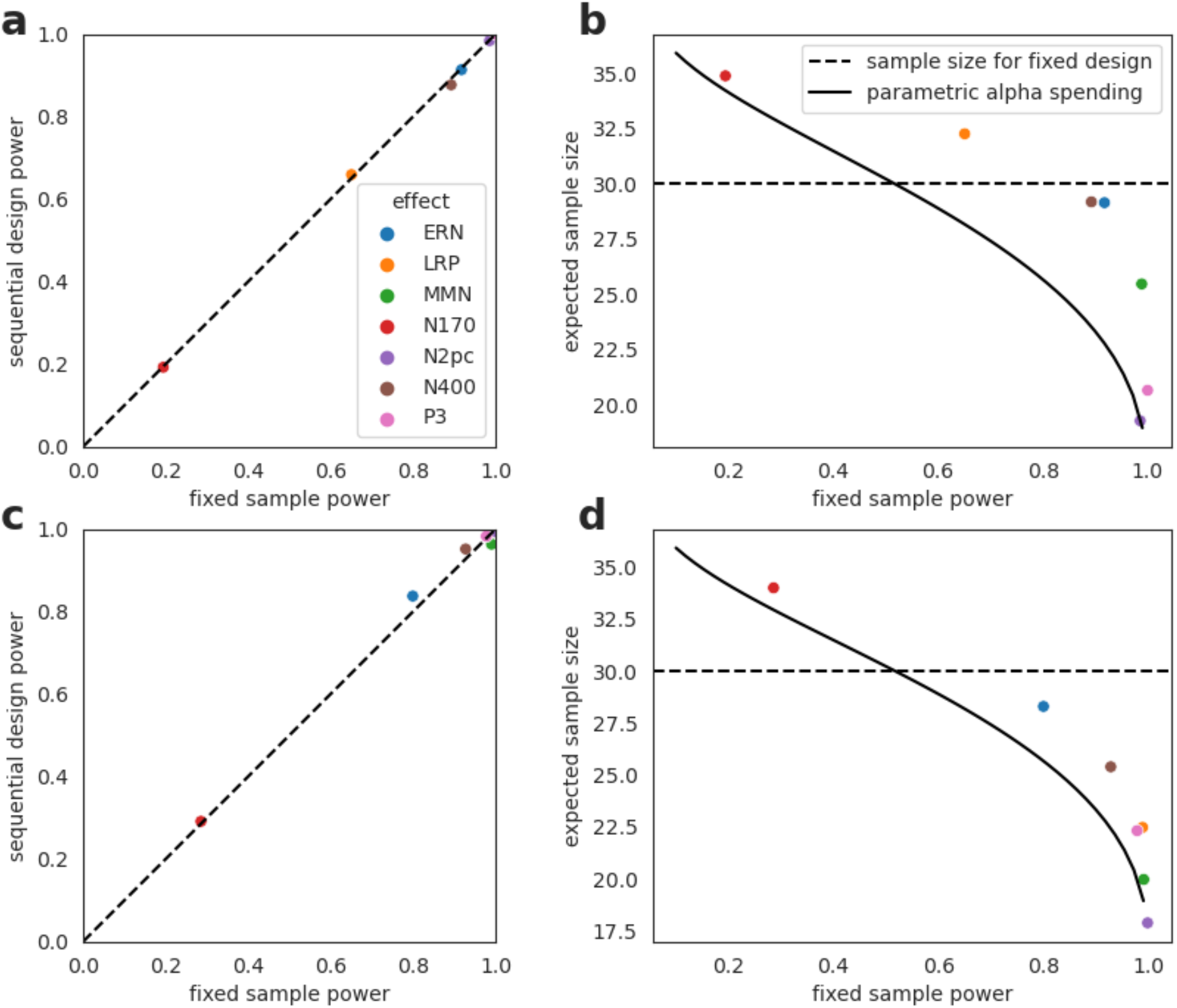
Results of power simulations. (a and c) Power for detecting ERP effects compared between fixed-sample designs with *n*_fixed_ = 30 and sequential designs with one interim analysis, a Pocock spending function, and *n*_max_ = 30 × IF, where IF is an inflation factor known a priori (see Section 2.1.2). **(b and d)** The expected sample size of the sequential designs as a function of statistical power for detecting the effect of interest, compared to the corresponding sample-size vs. power curve for parametric sequential designs, which may be considered a lower limit for expected sample size. (**a) and (b)** show the results for a *t-*max test, while **(c) and (d)** show the results for a cluster-based permutation test.

When the study design is well-powered for detecting an effect, the expected sample size is smaller because the probability of rejecting the null hypothesis at an interim analysis is higher Consequently, for sufficiently well-powered designs, we see up to >30% sample size saving compared to a fixed-sample design with the same power (see Figure 3b and 3d). Conversely, if power is very low, then the study will often run to completion without detecting the effect and the expected sample size will be close to the maximum sample size, which is greater than the fixed sample size. While we do see efficiency gains over fixed-sample designs, we find that permutation alpha spending is not as efficient as parametric alpha spending (see Figure 3b and 2d); this is to be expected, as parametric methods are generally more efficient than non-parametric methods when their assumptions are met. For the statistical tests used in these simulations (*t-*max and cluster-based permutation tests), the assumptions of parametric alpha spending are *not* met, but we show the theoretical curve as a lower bound on the expected sample sizes we could in-principle anticipate from our permutation approach. Interestingly, cluster-based permutation tests are closer to the lower bound than is the *t-*max procedure, so the efficiency gains one can expect from permutation alpha spending may depend on the statistical test used, in contrast to parametric alpha spending (where relative efficiency depends only on the design parameters, e.g. spending function and number of interim analyses). However, gains should always be seen when the probability of early rejection is high.

## 4. Discussion

### 4.1 Use cases of sequential testing for neuroimaging

We anticipate at least several main use cases for permutation alpha spending in human neuroscience.

- A researcher does not have any reasonable means of running an a priori power analysis. Instead, they select a conservative maximum sample size they would be willing to collect, but they use permutation alpha spending to perform interim analyses throughout data collection while controlling their false positive rate. If the effect size ends up being large enough to detect with a smaller sample size, they will likely be able to stop data collection early.
- A researcher has conducted a power analysis for a fixed-sample design, but they found that they need to collect a very large sample size, so they wish take advantage of the efficiency gains afforded by a sequential design. To this end, they use an inflation factor to convert the fixed-sample design for which they’ve already done a sample size calculation into similarly-powered sequential design with several interim analyses. Now, if they find there is already enough evidence to reject the null hypothesis at an interim analysis, they may stop data collection early.
- A researcher has no reasonable means of running an a priori power analysis. Instead, they select an initial maximum sample size, and they perform a *conditional power analysis* by bootstrap after the first interim analysis, which they use to adjust their maximum sample size midway through data collection so as to achieve some desired power. In this way, they get to conduct a power analysis on the basis of some internal pilot data, but if the pilot data alone provide enough evidence to reject the null hypothesis, they need not collect any additional data.

In the last example, the researcher could conduct a power analysis very similarly to how we have conducted the power analyses above. However, a *conditional* power analysis differs somewhat from an a priori power analysis in that it estimates the probability of a given design rejecting the null hypothesis *given the data already collected* (Spiegelhalter et al., 1986). We have included an (experimental) module for estimating conditional power by Bayesian bootstrap (Rubin, 1981) in our Python package, though only power analyses for one-sample (or paired-sample) tests are implemented at time of writing.

Also note that, if one adjusts the maximum sample size midway through, the alpha spending function must be adjusted accordingly to reflect the new design or the false positive rate will not be controlled (Lakens et al., 2021). One would similarly need to change the alpha spending function if, for example, they end up collecting fewer observations than their originally intended maximum sample size due to practical constraints. We provide an example of how to adjust one’s alpha spending function mid-experiment on our package’s GitHub and Zenodo repositories.

Even the researcher in the second example above may wish to run a conditional power analysis during their study if they estimated their sample size based on a previously published study, since that previous study may have overestimated its effect size as a result of low statistical power, publication bias, or selective reporting (e.g. reporting the effect sizes within significant clusters in a fMRI study) (Poldrack et al., 2017). The ability to adjust one’s design on the basis of a conditional power analysis affords more options for navigating biases in the published literature while designing their own, well-powered study.

### 4.2 Stopping for futility

In the simulations we feature in this article, we assume that data collection continues until the specified maximum sample size unless the null hypothesis is rejected at an interim analysis.

Another option, however, is to run a conditional power analysis after the first (or any/each) interim analysis to estimate the probability of rejecting the null hypothesis if the design is run to completion. If this power falls below some (predetermined) threshold, then one might choose to stop the study for futility, rather than waste resources by continuing (K. K. G. Lan & Trost, 1997). In this case, one might achieve even greater sample size savings than we have described above.

However, it is important to note that including a stopping rule for futility in one’s design may affect the statistical power of the design. See work by Lakens and colleagues for a somewhat more thorough discussion (Albers & Lakens, 2018; Lakens, 2014; Lakens et al., 2021; Lakens & Evers, 2014). In principle, it is possible to estimate this effect by simulation, but this would require substantial computation (since power analyses would be nested within a larger power analysis, and the above simulations already took a great deal of compute time).

### 4.3 Reporting the results of a sequential analysis

There are not yet uniformly agreed-upon standards for reporting the results of sequential analyses. However, sequential analyses should report, minimally, the spending function used, the time of all interim analyses performed, the adjusted significance thresholds and value of the alpha spending function at each analysis, and the *p*-values observed at each analysis. In the case of *t-*max and other max-type procedure, as well as with threshold-free cluster enhancement, in which each voxel is assigned a *p*-value, we suggest reporting the smallest *p*-value obtained at each interim analysis and reporting full results for the time at which data collection was stopped. See the parametric alpha spending tutorial by Lakens and colleagues for more discussion (Lakens et al., 2021).

Note that it has been suggested that sequential analyses should be pre-registered for the sake of transparency (Lakens, 2014), which may be helpful in providing evidence that one did not change their design parameters in the analysis stage. Since the number and timing of interim analyses does not need to be determined a priori for the alpha spending procedure to control the false positive rate, then it is sufficient to specify a statistical test, an alpha spending function, a maximum sample size, and whether a stopping rule for futility will be used. However, it is also helpful to specify tentative interim analysis times, even though these may be altered without issue later on.

### 4.4 Limitations of the present approach

The present approach has a number of limitations that are worth discussing.

If one stops data collection for a neuroimaging study as soon as one accrues enough evidence to reject the null hypothesis on the basis of a cluster (e.g. in a cluster-level test) or a voxel (e.g. in a *t-*max procedure), one might miss other, more weakly activated clusters or voxels elsewhere in the data. In principle, this is also a limitation of fixed-sample designs with low statistical power, so a well-powered sequential design, we think, is still usually preferable to arbitrary or heuristic sample size determination.

Further, when a sequential design stops at an interim analysis, there is a risk that the test statistic crossed the rejection threshold because, due to random variation, the effect size was overestimated at that interim sample size. This does not inflate the false positive rate, as this risk is accounted for in the null distribution, but may result in biased effect size estimates for sequential designs. This may, however, be less of a problem for permutation alpha spending than for parametric alpha spending, because the exact numerical value of the test statistic used in the permutation test (e.g. cluster mass) is usually not directly of interest in neuroimaging studies. Moreover, this bias is washed out in meta-analyses, since effect sizes measured in studies that are terminated early are balanced out by those that ran to completion (Schönbrodt et al., 2017). This fact underscores the importance of sharing unthresholded statistical maps from neuroimaging studies, which can be hosted on platforms such as NeuroVault to facilitate future metanalyses (Gorgolewski et al., 2016).

Both of the above issues can also be optionally alleviated, if a researcher wishes, by continuing to collect data past the interim analysis at which the null hypothesis is rejected. There is nothing stopping data collection after the null hypothesis has already been rejected in the interest of yielding better estimates of the effect of interest – or, better yet, estimating that effect in an independent sample.

Finally, with any permutation-based method, one requires a sufficiently large sample size for valid inference. For instance, the number of possible permutations for a one-sample permutation test with *n* = 5 observations is 2^5^ = 32, which is far too small; the lowest p-value one could possibly compute with that few permutations is 1/32 = 0.031. Thus, an interim analysis at *n* = 5 has no chance of rejecting the null hypothesis if the adjusted significance threshold for that analysis is lower than 0.031 and still a substantially reduced chance otherwise. Researchers should ensure that the number of possible permutations at their smallest interim sample size is sufficient to reject the null hypothesis at the adjusted significance level determined by their alpha spending function.

### 4.5 Outlook

We believe that sequential designs can be a valuable tool as cognitive neuroscientists continue their efforts to improve the statistical power of neuroimaging studies, while balancing costs. Sequential designs provide multiple alternative paths to principled sample size determination in the event that conventional, a priori power analyses are difficult to perform. Moreover, even in the event that one can easily perform a power analysis for a well-specified effect of interest, highly-powered studies (e.g. confirmatory trials) stand to benefit greatly from the efficiency advantages of sequential designs. Indeed, when each subject costs hundreds or even thousands of dollars to run, as in an fMRI study, a greater than 30% reduction in expected sample size without sacrificing statistical power (see Figure 3) can free up valuable resources for cognitive neuroimaging labs and their funding agencies. We hope that our permutation-based approach to sequential analysis proposed in this article, and our accompanying Python package, empower cognitive neuroscience researchers to conduct more efficient studies.

## Acknowledgements

J.P.V. was supported by NSF GRFP DGE 1746045 and a Neubauer Family Distinguished Doctoral Fellowship; H.C.N. was supported by NSF NCS 2024923 and NSF NCS 1835181. This work was completed in part with resources provided by the University of Chicago’s Research Computing Center.

## Notes

https://github.com/john-veillette/niseq

https://doi.org/10.5281/zenodo.7517285

https://doi.org/10.5281/zenodo.7666443

### Competing Interest Statement

The authors have declared no competing interest.

